# Joint effects of balancing selection and population bottlenecks on the evolution of a regulatory region of human anti-viral *APOBEC3*

**DOI:** 10.1101/2023.07.18.549600

**Authors:** Naoko T Fujito, Sundaramoorthy Revathidevi, Yoko Satta, Ituro Inoue

## Abstract

Human anti-viral APOBEC3s are crucial components of the innate immune response, playing a key role in inhibiting viral replication and proliferation by inducing mutations in viral genomes. In this study, in a regulatory region of ∼16 kb in the *APOBEC3* gene cluster on human chromosome 22, we identified three distinct haplogroups that are maintained at high frequencies in both African and non-African populations today. Despite the long persistence of these haplogroups, one of which is shared by archaic hominins, the nucleotide diversity within each haplogroup was unusually low in non-Africans. The simultaneous observations of low intra-haplogroup diversity and high inter-haplogroup divergence were incompatible with neutrality, as demonstrated by the genome-wide empirical distribution and coalescence-based simulation tests. In addition to the three shared haplogroups, an ancestral sequence group was identified exclusively in African populations. To explain these features and investigate the underlying evolutionary mechanisms, we performed forward simulations to model the joint effects of balancing selection and population bottlenecks. We demonstrated the operation of balancing selection over the past ∼1 million years, as well as a hitherto unexplored reduction in intra-haplogroup diversity. The only unexplainable observation was the low diversity within African haplogroup I, which is unaffected by the non-African-specific bottleneck. We hypothesized that an extra haplotype turnover within haplogroup I occurred prior to the Out-of-Africa dispersal of modern humans. Finally, we discussed the functional importance of the *APOBEC3* regulatory region in terms of evolutionary conservation in Hominidae, a target of natural selection in modern humans, and its relevance in GTEx eQTL.

**Significance statement:** This study provides new insights into the joint effects of population bottlenecks and balancing selection on genetic diversity within haplogroups. First, we reveal how balancing selection interacts with demographic history to shape intra-haplogroup diversity, a previously unexplored aspect. Second, our findings are generalizable, as illustrated by the ABO locus—an established example of balancing selection—where low within-haplogroup diversity and high between-haplogroup divergence are also observed in non-African populations. Third, the involvement of balancing selection in a highly conserved regulatory region suggests functional significance in modulating the expression of all seven APOBEC3 genes. While further validation is needed, our findings underscore the evolutionary and functional importance of this region in human innate immunity.

## Introduction

APOBEC3 belongs to a family of cytidine deaminases that play essential roles in the innate immune response against retroviruses, endogenous retroelements, and DNA viruses. They catalyze the deamination of cytidine in single-stranded DNA or RNA, converting cytidine-to-uridine (C > U). This C-to-U hypermutation disrupts the genomic information of viruses, inhibiting their replication and proliferation and resulting in their degradation via the uracil DNA glycosylase (UNG2)-dependent pathway. APOBEC3s also inhibit viral replication through deaminase-independent mechanisms (Newman et al. 2005; Wang et al. 2012; Pollpeter et al. 2018; Hakata et al. 2019).

The APOBEC3 family consists of seven enzymes (APOBEC3A/B/C/DE/F/G/H), and their genes are clustered within a 200-kb region on chromosome 22 (the *APOBEC3* gene cluster, Figure 1a). Although each member exhibits distinct characteristics, including cellular localization, tissue-specific expression, substrate specificity (e.g., single-stranded DNA or RNA), and viral restriction properties, their functions show mild overlap. For example, while APOBEC3G acts as the most potent anti-human immunodeficiency virus (HIV) factor, APOBEC3D, 3F, and 3H also contribute to HIV restriction. Similarly, APOBEC3A, 3B, 3C, 3F, and 3G can restrict DNA viruses, including hepatitis B virus, human papillomavirus, adeno-associated virus, herpes simplex-1, and Epstein-Barr virus (Stavrou and Ross 2015; Revathidevi et al. 2021).

**Figure 1.**
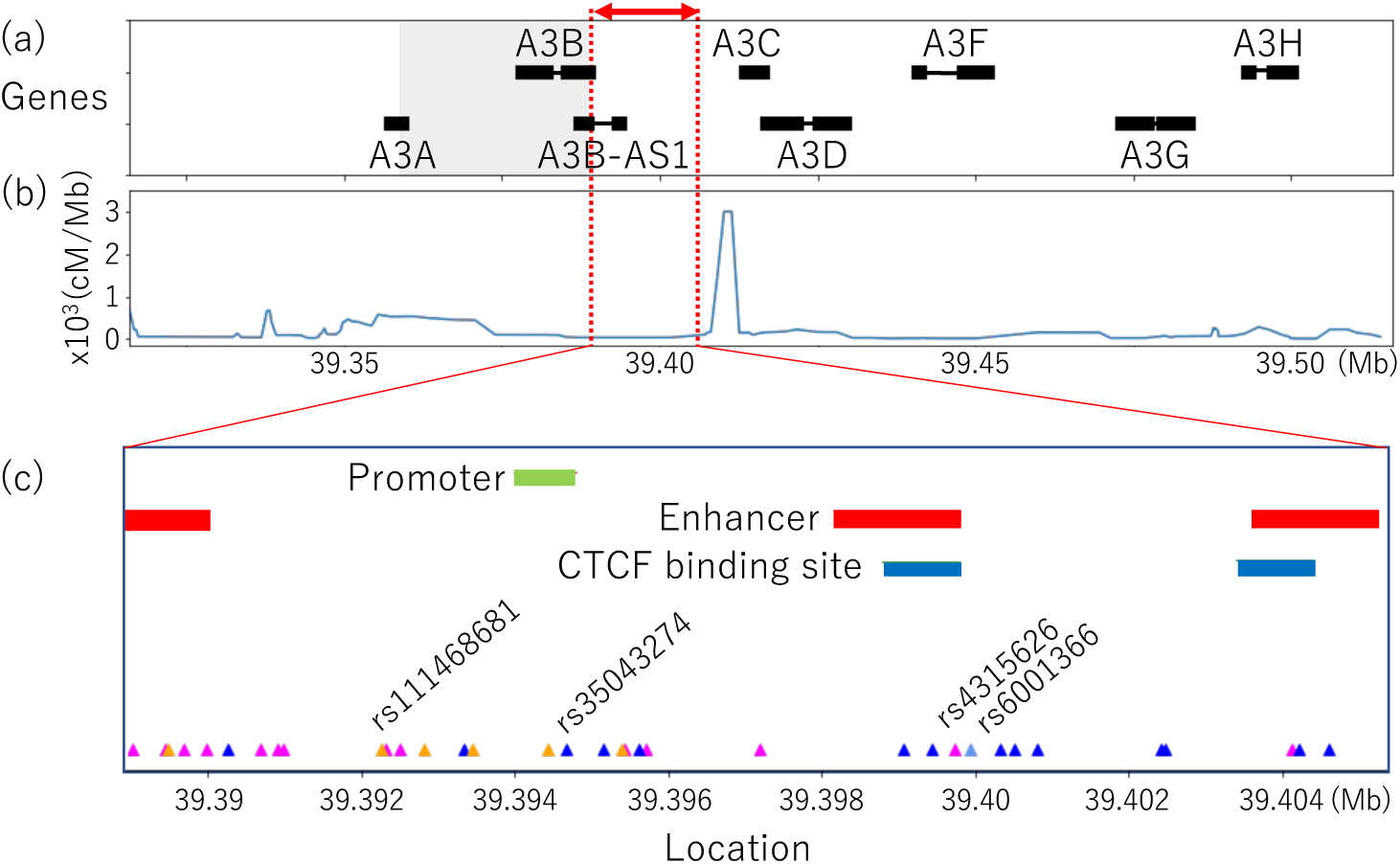
The APOBEC3 cluster including the regulatory region. (a). The APOBEC3 cluster region, which spans 200 kb surrounding the APOBEC3 genes, is illustrated. A3A, A3B, A3C, A3D, A3F, A3G, and A3H are abbreviations for APOBEC3A, APOBEC3B, APOBEC3C, APOBEC3D, APOBEC3F, APOBEC3G, and APOBEC3H, respectively. A long non-coding RNA, APOBEC3B antisense RNA 1 (A3B-AS1), is also present in this region. The 16-kb regulatory region is represented by the double-headed red arrow, and the APOBEC3B deletion polymorphism is indicated by the shaded square. (b) The recombination rate in the APOBEC3 cluster region is plotted based on the HapMap phase II genetic map. (c) Annotated regulatory sequences in the Ensemble Regulatory Build. The SNPs in LD (r^2^> 0.8) with the index SNPs for HI (magenta), HII (blue), and HIII (orange) are depicted with triangles.

In the present study, while initially analyzing patterns of genetic variation around the *APOBEC3* gene cluster, we noted an approximately 16-kb long non-coding region within the cluster exhibits three distinct haplogroups in all 26 populations in the 1000 Genome Project (TGP), each of which is strongly supported by a long branch in the genealogical tree. Although this region does not contain any coding sequences, it harbors several regulatory motifs. Its functional importance is evidenced by evolutionary conservation across the Hominidae family. Interestingly, each haplogroup in this region displays a star-like burst of closely related lineages. We refer to this pattern as multi-star genealogy and examined the frequency of this observation in the results of coalescence simulation under neutrality, as well as in the empirical distribution obtained through a genome scan. The results from both the simulations and empirical analyses were incompatible with the observed multi-star genealogy, prompting us to reject the null hypothesis of neutrality and to consider an alternative explanation. In doing so, we further identified one important difference between African and non-African genealogical trees: the presence of apparent “ancestral” haplotypes that occur at appreciable frequencies in African populations but are almost absent in non-African populations.

Our alternative hypothesis assumes balancing selection that can maintain multiple alleles in a population for a long time. Two well-known forms of balancing selection are negative frequency-dependent selection and overdominance selection, although the patterns of genetic variability yielded by those mechanisms are often indistinguishable from each other (Takahata and Nei 1990; Charlesworth 2006 and Fijarczyk et al. 2015 for reviews). In recent years, with the advent of next-generation sequencing, numerous studies have rigorously identified candidate loci undergoing balancing selection. There is a consensus that balancing selection primarily affects a limited category of genes, with disease-resistance genes being the most well-known example (Bitarello et al. 2023 for a review). Balancing selection can extend the most recent common ancestor (MRCA) of each allelic group further back in time than genetic drift alone, and it may occasionally result in trans-species polymorphism as observed at major histocompatibility complex (MHC) loci (Takahata 1990; Takahata and Nei 1990). However, as almost all studies on genetic variation under balancing selection have primarily focused on nucleotide divergence among different allelic groups, nucleotide diversity within an allelic group is largely overlooked. Therefore, we performed forward simulations to explore the joint effects of balancing selection and population bottlenecks.

We also discussed the functional differences among the three haplogroups by analyzing the distribution of derived alleles within the regulatory region, associating them with Genome-Wide Association Study (GWAS) hits linked to these haplogroups, and incorporating expression Quantitative Trait Loci (eQTL) analyses from the GTEx database. Our results provide insights into the functional importance of the *APOBEC3* regulatory region in human evolution during the Upper Pleistocene.

## Results

### Non-African and African haplogroups in the regulatory region

The 16-kb non-coding region is located between *APOBEC3B* and *APOBEC3C*, flanked by a recombination hotspot and a deletion polymorphism within *APOBEC3B* (Figures 1a, 1b, S1 and S2). We term this region the *APOBEC3* regulatory region as it harbors one promoter, three enhancers, and two CTCF-binding sites, as annotated by the Ensemble Regulatory Build (Figure 1c). When we constructed genealogical trees of haplotypes within this region, it became immediately apparent that in any non-African population in the TGP, three distinct groups of haplotypes (haplogroups, Figure s2a–d and S3) were present. These haplogroups exhibited appreciable frequency (*x* > 0.15), extremely low nucleotide diversity within each haplogroup (π < 0.01%), and relatively large nucleotide divergence between a pair of haplogroups (*d* >0.01%) (Tables 1 and S1). Using the minimum *x*, the smallest *d*, and the highest π values observed among the 15 non-African populations, we defined the three haplogroups I, II, and III (hereafter denoted as HI, HII, and HII, respectively; see Materials and Methods). The divergence time between HI and HII was estimated to be as old as 1.80–1.96 million years ago (MYA), assuming that the *de novo* mutation rate in humans and hominoids is 0.5 × 10^-9^ site/year (Jónsson et al. 2017; Chintalapati and Moorjani 2020), whereas those between HI and HIII and between HII and HIII were estimated to be 1.36–1.40 MYA and 1.43–1.55 MYA, respectively (Table S1). In contrast to these long divergence times, intra-haplogroup diversity is one order of magnitude lower than expected when calculated by multiplying the nucleotide diversity in all pairwise comparisons by the haplogroup frequency (Tables 1 and S1). We term this unique structure, characterized by a genealogical tree exhibiting multiple star-like bursts, “multi-star” genealogy. Hereafter, we mainly aimed to elucidate the evolutionary history and mechanisms reflected in this unprecedented genealogical configuration.

To gain insight into the origins of the three haplogroups, we examined their presence in the African populations. All the African populations in the TGP possessed HI, HII, and HIII. However, among the African haplogroups, only HI exhibited extremely low intra-haplogroup diversity, whereas HII and HIII displayed moderate levels of diversity that are consistent with the expectation (Tables 1 and S1). In addition, over 40% of the haplotypes in Africans were unique to their populations and did not belong to any of the three haplogroups (Figure 2e, f). The genealogical tree of this genomic region in African populations indicated that the three haplogroups emerged independently from these African-specific sequences roughly 1 MYA. This finding suggests that the African-specific haplotypes are collectively an ancestral type (Figures S3c and S4, Table S2b).

**Figure 2.**
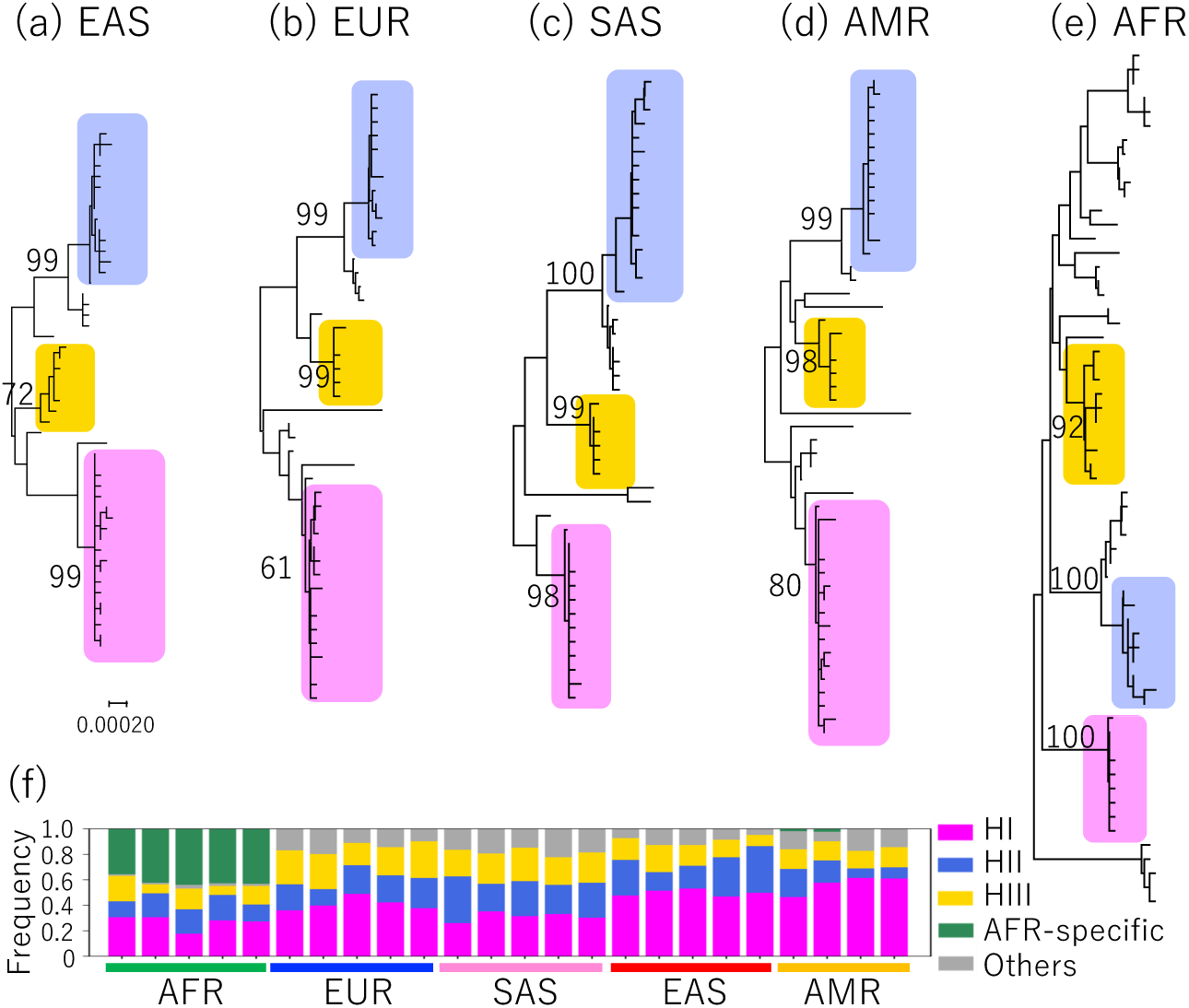
The three haplogroups observed in the APOBEC3 regulatory region. Phylogenetic trees were constructed using the Neighbor-joining method based on haplotype sequences identified in East Asian (a, 49 haplotype sequences from JPT), European (b, 45 haplotype sequences from CEU), South Asian (c, 45 haplotype sequences from PJL), American (d, 50 haplotype sequences from PUR), and African (e, 61 haplotype sequences from LWK) populations. Bootstrap values after 1000 replications are presented only for the three haplogroups: HI (magenta), HII (blue), and HIII (orange). The trees are drawn to scale, with branch lengths in the same units as those of the evolutionary distances used to infer the phylogenetic tree. The evolutionary distances were computed using the Maximum Composite Likelihood method and are displayed in the unit of base substitutions per site. (f) Proportion of the three haplogroups in each population in the 1000 Genomes project data. The sequences that do not belong to any of the three haplogroups are separated into two groups: African-specific sequences (green) and others (gray).

Although the estimates of the above emergence time of the haplogroups are necessarily approximate, their antiquity is supported by genomic data from ancient hominins. Specifically, all four pseudo-haploid sequences from the Altai Neanderthal (Prüfer et al. 2014) and Denisovan genomes (Meyer et al. 2012) used in this study harbor HI-linked alleles at three out of 14 SNP sites. These derived alleles are shared by all HI sequences but not by HII or HIII. Although the genealogical clustering of HI with the ancient hominin sequences is only weakly supported by bootstrap probabilities (Figures S3 and S4a, Table S2a), it is suggested that when archaic hominins and modern humans diverged (550 KYA; Prüfer et al., 2014), the HI haplogroup had already been isolated from other haplotypes.

The functional differences among the three haplogroups remain unknown. However, a window-based analysis of nucleotide differences revealed that regions with the highest divergence coincided with regulatory sequences, such as promoters and enhancers (Figure S5). This finding suggests potential functional differences among the haplogroups.

To investigate changes in the frequencies of the three haplogroups over time, we utilized genome-wide data from prehistoric humans. To this end, we analyzed the Allen Ancient Data Resource (Anon) for index SNPs for the three haplogroups and found that rs2142833 (*r*^2^ = 0.92 with HI in the global population), rs9607601 (*r*^2^ = 1 with HII in the global population, *r*^2^ = 0.87 in the JPT, and *r*^2^ = 0.55 in the CEU), and rs11089916 (*r*^2^ = 1with HIII in CEU and JPT) were informative. Due to the limited number of available ancient samples, we divided the data into different time windows of varying sizes (400**–**20,000 years). Although we were unable to analyze African data due to an insufficient sample size, we observed the three haplogroups have been maintained at similar frequencies with only moderate changes in both Europe and East Asia during the past 40,000 years (Figure 3).

**Figure 3.**
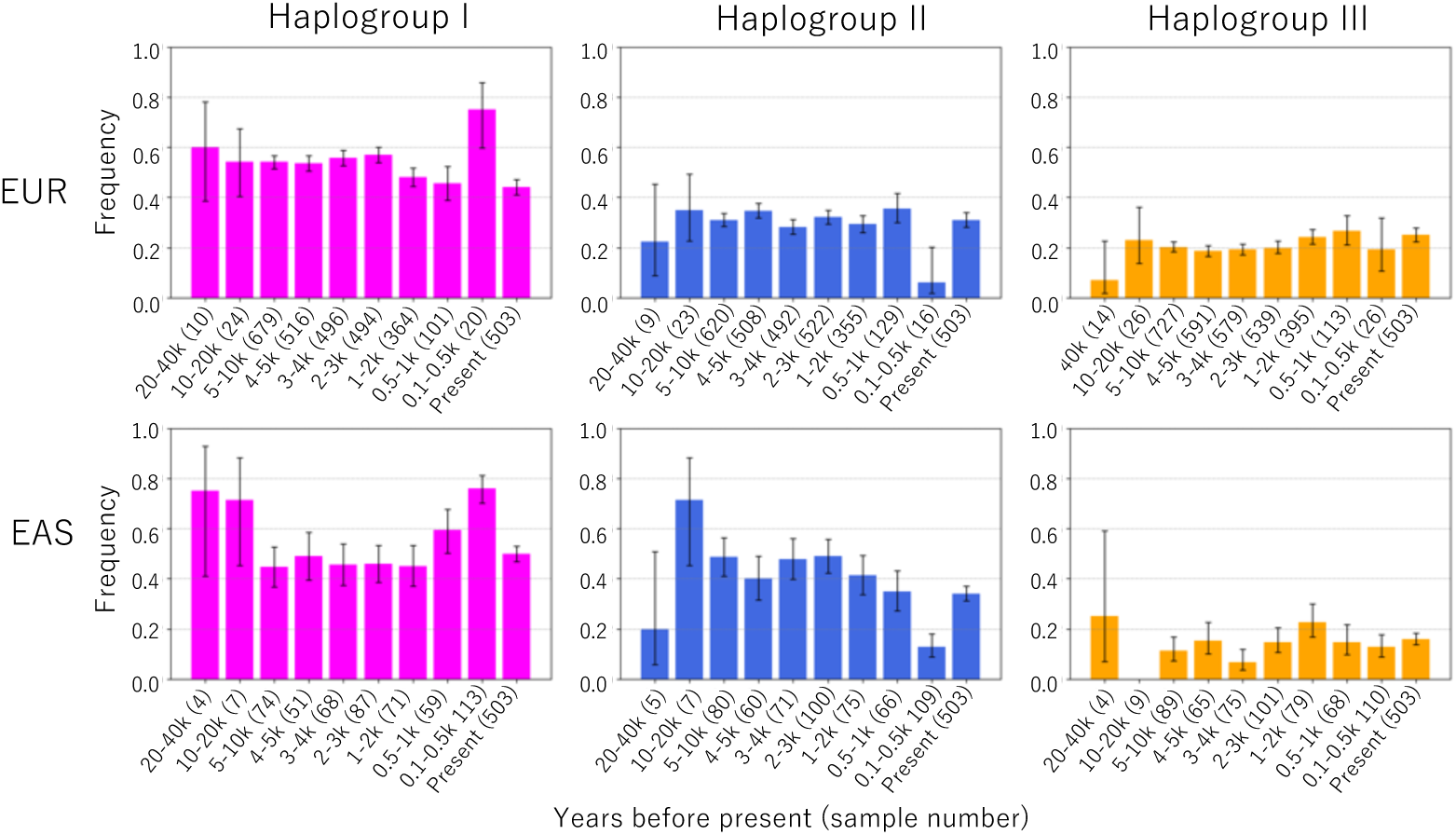
Frequencies of the three haplogroups in modern humans across time. Samples from Europe (top) and East Asia (bottom) are from compiled ancient data from the Allen Ancient Genome Resource and current data from the 1000 Genome Project. Frequency of the allele linked to each haplogroup was calculated at the informative SNPs (rs2142833, rs9607601, and rs11089916 for HI, HII, and HIII, respectively). As frequency was calculated after removing the samples for which genotypes were not available at each of the informative SNPs, the sum of the frequencies of the three informative SNPs is not necessarily equal to 1. The number of samples is shown in parentheses. Time periods are indicated in years before the present. Error bars indicate Wilson’s confidence interval (95%).

### Neutrality test for the regulatory haplogroups

The long-term persistence of the three haplogroups suggests the possibility of non-neutral evolution of the *APOBEC3* regulatory region. We tested the neutrality of the multi-star genealogy using coalescence simulation and found its extreme rarity (Materials and Methods). More specifically, we assessed the statistical rarity of the multi-star genealogy in African and non-African populations by evaluating (1) the frequency of occurrence of three haplogroups with sizes (i.e., the number of chromosomes) comparable to those of HI, HII, and HIII, and (2) the likelihood of co-occurrence of low intra-haplogroup diversity and high inter-haplogroup divergence. First, we searched for a genealogical structure with three distinct haplogroups, each having a size equal to or larger than the observed one. We conducted neutrality simulations by placing a specified (observed) number of segregating sites on a coalescent tree using ms (Hudson 2002). We used two different demographic models. The first is that proposed by Schaffner et al. (2005), which includes the Late Pleistocene Out-of-Africa bottleneck. The second is a modified version of the SFD model, which includes the bottleneck that occurred in Africa immediately before the Chibanian period, around 0.9 MYA, as recently inferred by Hu et al. (2023). It should be noted that the latter bottleneck effect cannot be completely erased by the subsequent population expansion with *N_e_* = 20,000 (Figure 3), approximately 30,000 generations between the two bottlenecks are not long enough for nucleotide diversity to equilibrate (Charlesworth and Jensen 2021). Of the 10^6^ coalescence trees we generated, 6159 (SFD model, *p =*0.62%) or 7628 (HFD model, *p =*0.76%) exhibited the presence of three distinct allelic groups of comparable sizes in JPT, where the haplogroups showed relatively high frequencies (Table S3; *x* = 50.0%, 36.5%, and 9.6% for HI, HII, and HIII, respectively). Such occurrence was more frequent when the haplogroup frequencies were low. For CEU, 42916 or 36858 out of 10^!^ replicates (*p* = 4.29% with the SFD model and *p* = 3.69% with the HFD model, respectively) exhibited the presence of three distinct allelic groups with the required sizes (Table S3; *x* = 41.9%, 21.2%, and 22.2% for HI, HII, and HIII, respectively). These *p*-values suggest that the likelihood of multi-star genealogy is moderately rare in non-African populations, whereas the three haplogroups with smaller sizes occur rather frequently in African populations (Table S3).

Considering that three haplogroups are present, we investigated how often the observed levels of low intra-haplogroup diversity and high inter-haplogroup divergence occur. Using the coalescence simulation again, we collected 10^4^ replications where three haplogroups are present with the same sizes as in HI, HII, and HIII. We compared the observed nucleotide differences within and between the haplogroups (Table S1) to simulated ones to evaluate statistical significance. The comparison revealed that only a small proportion of the 10^4^ simulation results yielded the observed levels of nucleotide differences, with *p* ≤ 9.1 × 10^-6^ for JTP and *p* ≤ 6.6 × 10^-5^ for CEU (Table S3). This finding suggests that some forms of natural selection have been involved in the emergence of the multi-star genealogy. It is worth emphasizing that the co-occurrence of three low-frequency haplogroups in Africans is not unlikely; however, intra-HI diversity is significantly reduced even in Africans, as in non-Africans (Supplementary Text). This results in *p* ≤ 1.2 × 10^-4^ for LWK and *p* ≤ 2.3 × 10^-4^ for YRI (Table S3).

We also examined genomic data for the multi-star genealogy using a window-based approach. We identified 2,460 regions (45,799 windows) in the JPT population that displayed three haplogroups equal to or larger than those observed, corresponding to 80,464 kb or 2.9 % of the total sequence analyzed (2.8 Gb). In the CEU population, we identified 13,507 genomic regions (298,169 windows of 489,448 kb length), accounting for 17.5% of the total sequence analyzed. These observations collectively imply that while the occurrence of the multi-star genealogy may be uncommon in human genomes, the likelihood strongly depends on the frequency of haplogroups. To examine the intra-haplogroup diversity, we focused on genomic regions in which each of the three haplogroups occurs with the same frequency as the observed one. We identified 260 and 344 such regions for JPT and CEU, respectively, none of which displayed the observed level of intra-haplogroup diversity.

In summary, the structure of genealogy with the three haplogroups in the *APOBEC3* regulatory region is rarely found in coalescence simulation or in actual genomic data. Even when we collected such structures from tens of millions of replications of coalescence simulations or from the whole genome, the likelihood of simultaneous occurrence of low intra-haplogroup diversity and high inter-haplogroup divergence remained exceedingly low. We thus conclude that these haplogroups have unlikely evolved in neutral fashions; rather, their long persistence suggests the involvement of some forms of balancing selection that has persisted in African and non-African populations over the past 1 million years.

### Balancing selection and population size changes

We then focused on differences in the patterns of genetic diversity between African and non-Africa populations. Although balancing selection was suggested for both populations, the pattern and level of intra-haplogroup diversity were evidently different between these two meta populations (Table 1). In non-African populations, the intra-haplogroup diversity was 10 times lower than expected. In contrast, African populations showed high and consistent levels of diversity within HII and HIII, in line with expectations. However, the intra-HI diversity exhibited a substantial reduction not only in non-African populations but also in African populations. Hence, these features require further scrutiny.

**Table 1.** Intra- and inter-haplogroup diversity and divergence of HI, HII, and HIII.

| Population | Haplogroup | Frequency | $\pi$ or $d$ (%) | All pairwise $\pi$ (%) |
| --- | --- | --- | --- | --- |
| JPT | HI | 0.50 | 0.0056 | 0.10 |
|  | HII | 0.37 | 0.0065 |  |
|  | HIII | 0.09 | 0.0096 |  |
|  | HI - HII | - | 0.1821 |  |
|  | HI - HIII | - | 0.1394 |  |
|  | HII - HIII | - | 0.1448 |  |
| CEU | HI | 0.41 | 0.0073 | 0.10 |
|  | HII | 0.21 | 0.0062 |  |
|  | HIII | 0.22 | 0.0039 |  |
|  | HI - HII | - | 0.1803 |  |
|  | HI - HIII | - | 0.1386 |  |
|  | HII - HIII | - | 0.1482 |  |
| LWK | HI | 0.27 | 0.0028 | 0.14 |
|  | HII | 0.13 | 0.0198 |  |
|  | HIII | 0.15 | 0.0168 |  |
|  | HI - HII | - | 0.1894 |  |
|  | HI - HIII | - | 0.1361 |  |
|  | HII - HIII | - | 0.1529 |  |

Intra-haplogroup diversity ( $\pi$ ) and inter-haplogroup divergence ( $d$ ) for each haplogroup in JPT, CEU, and LWK populations are presented together with haplogroup frequency and overall diversity in the *APOBEC3* regulatory region (all pairwise $\pi$ ).

To explain these differences between African and non-African populations, we developed a forward simulation model that incorporates balancing selection and population size changes with one or two bottlenecks. In this model, we considered four different types of sequences: three representing the three haplogroups and one additional group, hereafter referred to as “ancestral group”, representing the African-specific sequences. Given the potential roles of the haplogroups in regulating the expression of the anti-viral APOBEC3s, we first considered negative frequency-dependent selection and hypothesized a scenario in which the ancestral group played a fundamental role in gene regulation, and the new haplogroups that subsequently emerged one after another modified this original regulatory function independently. These functions might include enhancing the expression of certain *APOBEC3* genes or silencing others, thereby conferring robustness against specific viral groups. We thus assumed that each haplogroup is subject to negative frequency-dependent selection with their relative fitness given by 1 − *sx*, where *x* is the frequency of a haplogroup and *s* is selection coefficient common to the three haplogroups. In contrast, the relative fitness of the ancestral group is defined as 1 − *as*, where *a* is a constant parameter. In our model, the ancestral group is not subject to balancing selection because the actual African-specific sequences are not monomorphic and do not share a common mutation upon which balancing selection could act. The fitness for diploids is assumed to be multiplicative.

We used demographic models similar to those used for the coalescence simulation (Figure 4a, b). This time, we included a recent population expansion, though it does not significantly affect genetic diversity within and between sequence groups (Figures 4e-h, S6, and S7). For simplicity, all cases assumed that three haplogroups and an ancestral group emerged simultaneously, with the initial frequency of 0.25 set prior to the pre-Chibanian bottleneck, and are immediately subjected to frequency-dependent selection. Simulations were performed with a range of parameters *s* and *a*, and we examined intra- and inter-haplogroup diversity together with frequency changes of the three haplogroups and the ancestral group. Here and thereafter, the value of *s* is scaled relative to *N*_*e*_ specified in the simulations.

**Figure 4.**
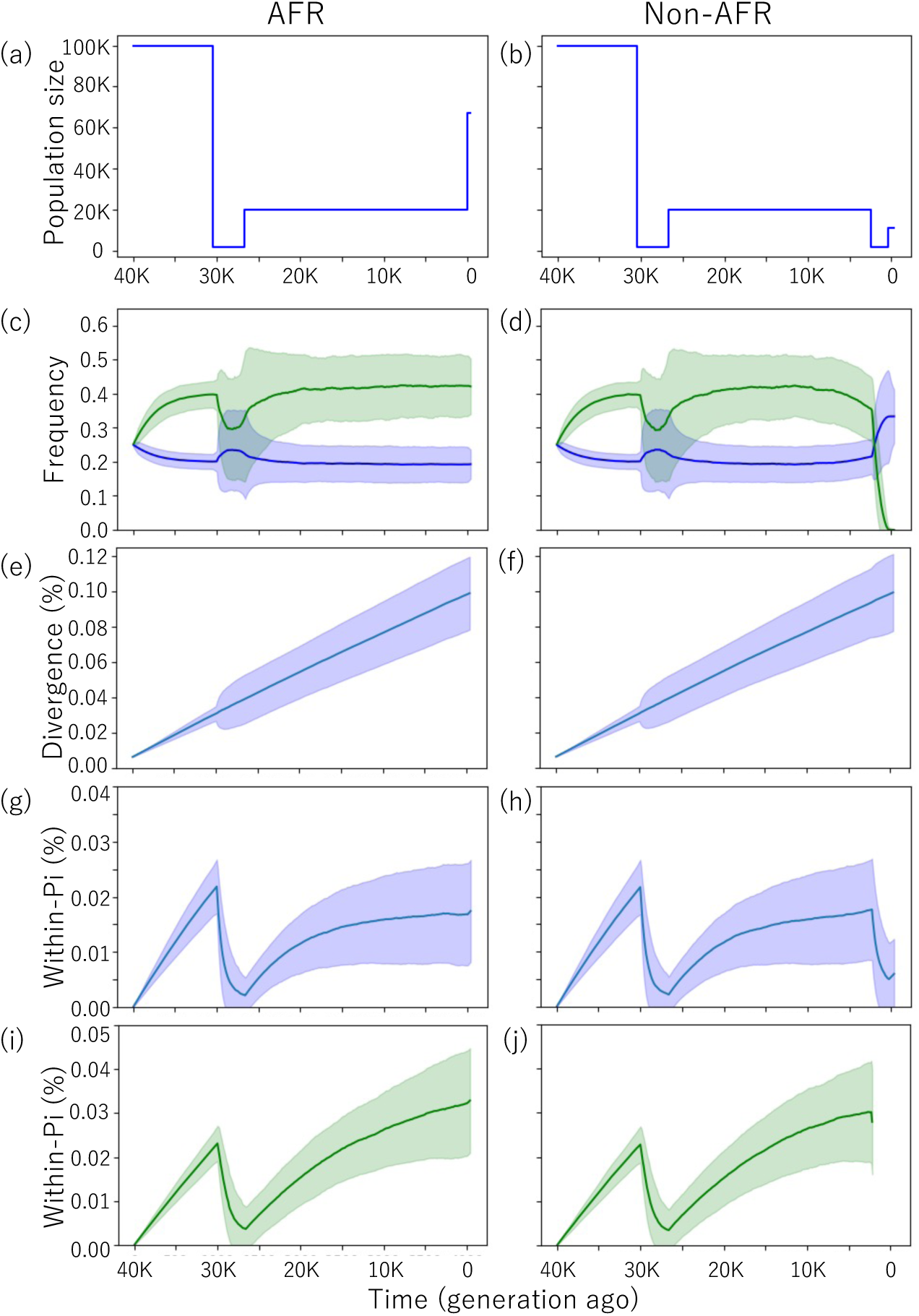
Simulation study using a model of negative frequency-dependent selection with demographic models for African (a) and non-African populations (b). The simulations assumed three haplogroups and one ancestral group, with relative fitness for each haplogroup given by 1 − *sx*. The ancestral group fitness was assumed to be 1 − *as*. The mutation rate was assumed to be 1.25 × 10^-8^ per site per generation. Simulations were conducted with population sizes and time duration reduced by a factor of 1/10 and a mutation rate increased by a factor of 10. (For the initial phase, the mutation rate was further increased by a factor of 5 to reduce the computational time.) Shifts in haplogroup frequency, intra-haplogroup diversity, and inter-haplogroup divergence are illustrated for conditions with *s* = 0.05, *a* = 0.2 for Africans and non-Africans. Only replications in which the observed state was replicated (i.e., the persistence of the three haplogroups and the ancestral group throughout in Africans) were utilized. For non-Africans, replications were considered in which the three haplogroups persisted, while the ancestral group was lost during the Out-of-Africa bottleneck. Lines and shaded areas represent the mean values and standard deviations across 1,000 replications in which the observed sequence groups persisted. Frequency trajectories for haplogroups and the ancestral group are depicted with blue and green lines, respectively, for African (c) and non-African populations (d). Shifts in simulated inter-haplogroup diversity are presented for African (e) and non-African (f) populations. Intra-haplogroup diversity is similarly presented for African (g) and non-African (h) populations. Diversity within the ancestral group was also shown for African (i) and non-African (j) populations.

The results demonstrated that intra-haplogroup diversity is not affected by the value of *s*, but sensitively varies depending on the value of *a* (Figures 4g, 4h, S6a and S7a). This variation arises simply from differences in haplogroup frequency (Figures 4c, 4d, S6b and S7b). In our model, the haplogroup frequency is a function of the constant parameter *a* (Supplementary Text). The larger the value, the lower the frequency of the ancestral group and the higher the frequency of the three haplogroups. In contrast to intra-haplogroup diversity, inter-haplogroup divergence grows linearly in time even for different sets of parameter values (Figures 4e, 4f, S6c and S7c), as long as selection is effective. When *a* = 0.2, the frequency reached the deterministic equilibrium value of 40 % for the ancestral group and 20 % for each haplogroup, as observed in African populations, showing that 0.2 is the proper value for *a* under negative-frequency dependent selection to explain the observation (Figure 4c, 4d, S6b and S7b). For African populations, the effect of the first bottleneck on intra-haplogroup diversity was somewhat faded at the end of the simulation (corresponding to ∼1 million years; Figure 4g). In the absence of the second bottleneck, intra-haplogroup diversity approached 0.02 % (Figures 4g and S6a), which is close to that observed for HII and HIII. However, as mentioned earlier, the observed diversity within HI is considerably small (Table 1) and cannot be explained solely by the present mutation scheme (Materials and Methods). The diversity within the ancestral group reached over 0.03 % (Figure 4i), whereas the observed diversity of the African-specific sequences was significantly higher, reaching 0.14 % and 0.15% in the combined African population (AFR) and the YRI population, respectively. This difference arises from the fact that the African-specific sequences are not monophyletic, and substantial genetic diversity had already accumulated prior to the onset of balancing selection. These values are nearly identical to the mean pairwise diversity within AFR and YRI overall, which is 0.15 % in both populations.

For non-African populations, the observed state is that all three haplogroups are retained, but the ancestral group is lost during the Out-of-Africa bottleneck. In the simulation, the ancestral group was more frequently lost during the first and more severe bottleneck (γ = 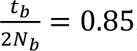, where *t_b_* and *N_b_* are the duration time in units of generations and the effective size of a bottleneck, respectively), but sometimes lost in the second, Out-of-Africa bottleneck (γ = 0.5). When we collected replications where the ancestral group was lost in the Out-of-Africa bottleneck, the intra-haplogroup diversity significantly decreased due to the bottleneck, reaching 0.005% (Figure 4h), which is close to that observed for HI, HII, and HIII in non-African populations. With the following recent population expansion, the intra-haplogroup diversity somewhat recovered, but the extent is small enough to be ignored (Figure 4h).

To assess whether a single combination of *s* and *a* values could account for the observations in both African and non-African populations, we conducted an additional simulation across a range of *s* and *a* values (Figure 5), using a demographic model that mimics the Out-of-Africa bottleneck (Figure 5a). The initial frequencies of the ancestral group and each haplogroup were 40 % and 20 %, respectively, as observed in African populations. All the sequences were immediately subjected to frequency-dependent selection and the Out-of-Africa bottleneck (Figure 5a, Materials and Methods). We calculated the proportion of simulation replications in which all the haplogroups were retained, but the ancestral group was lost, as observed in non-Africans. When examining the case with *a* = 0.2, the condition where *N*_&_*s* = 4 yielded the observed pattern at a rate as high as 5.3% (Figure 5b).

**Figure 5.**
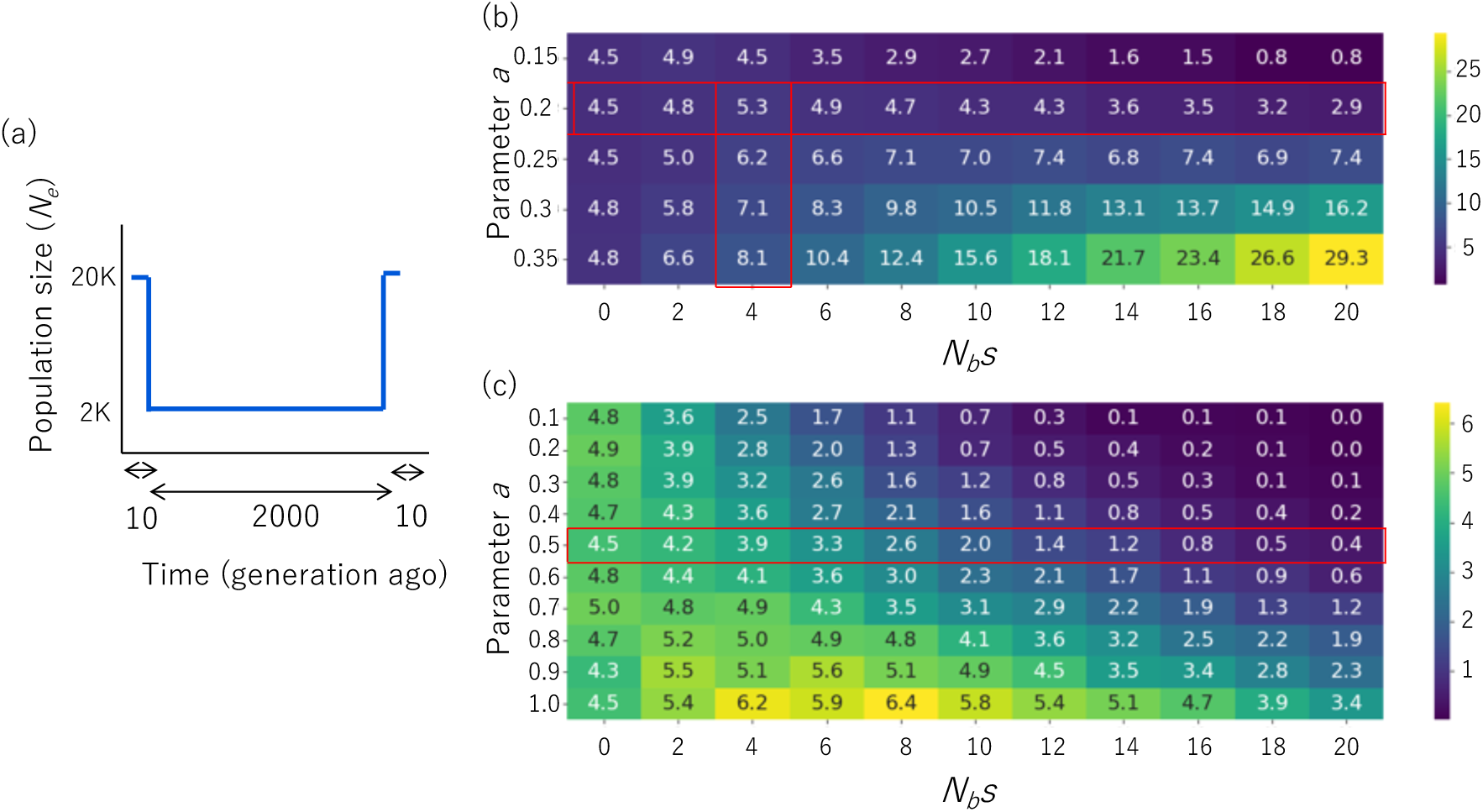
Probability of observed haplotype persistence during the Out-of-Africa bottleneck. The proportions of simulation replicates in which all three haplogroups persisted while the ancestral group was lost were calculated under models of negative frequency-dependent selection (b) and overdominance (c) each based on 10,000 replicates, and are presented as percentages. Panel (a) illustrates the demographic model incorporating a population bottleneck. The initial frequencies of the ancestral sequence group and each of the three haplogroups were 0.4 and 0.2, respectively. The red horizontal bars indicate the probabilities corresponding to the value of *a* that reproduces the initial frequencies under each model. The red vertical bars in (b) highlight the likely value of *N_b_s* that, in combination with the *a* value, produces the observed state with a probability greater than 5 %.

In summary, the extremely low intra-haplogroup diversity and high inter-haplogroup divergence observed in non-African populations can be explained by the combined effects of negative frequency-dependent selection and the Out-of-Africa bottleneck. The loss of the ancestral group in non-Africans and its retention in Africans can occur with the same set of *a* and *s* values (*a*=0.2 and *N_b_s* = 4). For African populations, reasonably high intra-haplogroup diversity of HII and HIII is explained by negative frequency-dependent selection acting similarly to population subdivision. However, the observed intra-haplogroup HI diversity cannot be realized from the present model.

To study other possible mechanisms of balancing selection, we examined overdominance or heterozygote advantage with some asymmetry in relative fitness. In our model, the relative fitness of homozygotes is given by 1 − *s* for the three haplogroups and 1 − *as* for the ancestral group. Although we used the same symbol *a* as in frequency-dependent selection model, the meaning is fundamentally different (Supplementary Text, Figures S8, S9). Under this overdominance model, the value of *a* that yields the equilibrium frequency of 40 % for the ancestral group and 20 % for each haplogroup was calculated to be 0.5 (Supplementary Text). In simulations of the Out-of-Africa bottleneck, we found that with *a* = 0.5, the observed state was reproduced in fewer than 5 % in 10,000 replications, regardless of values of *N_b_s* (Figure 5c). Although the difference in probability was small, these results suggest that balancing selection on the *APOBEC3* regulatory region is more likely to be driven by negative frequency-dependent selection than by overdominance.

### Turnover of haplogroup I

One possible explanation for the low intra-HI diversity in all populations is a haplotype turnover in HI. This haplogroup was estimated to have emerged around 2 million years ago, and its antiquity is further supported by the presence of HI-linked alleles in archaic hominins, as described above. Therefore, the low intra-haplogroup diversity observed in both African and non-African populations cannot be attributed to a recent origin of HI. In African populations, positive selection for HI can be detected by *F_c_* (Fujito et al. 2018). Other statistics, such as *G_c_* (in 2D SFS (Satta et al. 2019), suggest that the haplotype turnover began with a single HI haplotype (Table S4), resembling an ongoing hard sweep within the HI group. This turnover was not detected in non-African populations owing to the simultaneously reduced intra-HII and HIII diversity, mitigating the signal of positive selection on HI. We demonstrated in simulation that when a population bottleneck occurs shortly after the completion of a haplotype turnover, and there is insufficient time for the group carrying this haplotype to increase its intra-haplogroup diversity, the bottleneck does not substantially affect the diversity (Figure S10). This indicates that if the haplotype turnover within HI happened shortly before the Out-of-Africa bottleneck, then the intra-haplogroup diversity should be in a similar range in both African and non-African populations. Several observations support this conjecture. First, the estimated times of expansion of the haplogroups revealed that the TMRCA (time to MRCA) of both HII and HIII significantly differed between African and non-African populations. It is estimated between 30,000 and 80,000 years ago in non-Africans, considerably shorter than ∼350,000 years ago for HII and 180,000 years ago for HIII in Africans (Figure S11, Table S5). These recent TMRCAs in non-African populations are consistent with the Out-of-Africa bottleneck around 70,000 years ago. In contrast, the TMRCA of HI is between 50,000 and 80,000 years ago in both African and non-African populations. As the Out-of-Africa bottleneck cannot affect the TMRCA in African HI, the HI turnover is a likely alternative, although this does not necessarily imply independent operation of some forms of directional selection. Moreover, phylogenetic network analysis revealed that the most frequent HI haplotype appeared across Africans, Europeans, South Asians, East Asians, and Americans in TGP (Figures S12, Supplementary text). Thus, the present-day HI in the world population has expanded from this recent haplotype before African and non-African HI diverged from each other.

### Deletion polymorphism

The *APOBEC3* regulatory region is located adjacent to a large polymorphic deletion encompassing the entire coding region of *APOBEC3B*. This polymorphism has been repeatedly reported to increase cancer susceptibility (ICGC/TCGA Pan-Cancer Analysis of Whole Genomes Consortium 2020). To investigate whether balancing selection for the regulatory region was influenced by this deletion, we examined its evolutionary history. First, it is important to note that regardless of population, the deletion is predominantly linked to HI and only occasionally to HII, HIII, and the ancestral group (Figure S13), suggesting that it is of ancient single origin. Second, haplotypes with the deletion exhibit star-like genealogy only when it is linked to HI, and no such pattern is observed when linked to non-HI haplotypes (Figure S14a-e). Third, within HI, a large part of the most frequent haplotype carries the deletion in all five metapopulations in TGP (Figure S14). It was previously suspected that the deletion underwent positive selection, primarily based on its high frequency in Asia (∼35 % in East Asians and ∼15 % in South Asians) and its low frequency in Europe and Africa (∼6% and ∼0.9%, respectively). However, at that time, no clear conclusion could be drawn (Kidd et al. 2007). In this study, we demonstrated that Asian HI is subject to balancing selection and that the high frequency in Asia is simply due to the hitchhiking effect associated with the expansion of HI.

We further studied the possibility that the deletion may confer an additional selective advantage to haplotypes in HI. Equally advantageous sequences are obviously selectively equivalent within that selected group. On the other hand, if some sequences are more beneficial than others even in the same selected group, a nested selective sweep could be observed. We calculated *F_c_* values (Fujito et al. 2018) for the sequences within HI linked to the deletion in the Japanese population. The resulting *F_c_* value was not low (*p* = 0.49). Thus, in terms of selective advantage, the sequences linked to the deletion do not differ from other HI sequences. It appears that the deletion did not confer its own selective advantage and was, in fact, lost in Europe and Africa due to genetic drift.

## Discussion

Research on the diversity of alleles under balancing selection has primarily focused on inter-allelic or inter-haplogroup divergence, and, to the best of our knowledge, has paid little attention to intra-allelic or intra-haplogroup diversity. In the present study, our simulation results demonstrated that intra-haplogroup diversity under balancing selection greatly decreases when combined with population bottlenecks. In contrast, inter-haplogroup divergence is less affected by demographic changes, selection strength, haplogroup frequency, or detailed mechanisms of balancing selection. Instead, it is largely proportional to the time elapsed since haplogroups began to co-segregate under balancing selection.

The impact of an idealized instantaneous bottleneck can be evaluated using the parameter γ = *t_b_*/(2*N_b_*). Irrespective of the presence or absence of selection, *t*_&_ remains unchanged. However, the effective population size is influenced by balancing selection in a specific way: balancing selection acts as if the haplotypes are subdivided into *m* distinct lineages. This situation is analogous to population subdivision, such that under symmetric balancing selection, the effective size of each haplogroup may be reduced to *N*_&_/*m*. In other words, the extent of intra-haplogroup diversity maintained by balancing selection is subjected to an *m*-fold severe bottleneck compared to a non-subdivided scenario (*m*γ). A caveat, however, is that for large *m*, as in MHC, it becomes more likely that a certain number of haplogroups are lost due to genetic drift rather than that all haplogroups being retained and responding by reducing their intra-haplogroup diversity. This may still hold true even for *m* = 3. In our Out-of-Africa bottleneck simulations, all three lineages were not always maintained even under strong balancing selection.

Our results indicate that intra-haplogroup diversity maintained by balancing selection is lower in non-African populations than in African populations. To assess the generality of this pattern, we analyzed the ABO locus, a well-known example of balancing selection. As expected, the genealogical trees at the ABO locus exhibit lower intra-allelic diversity for the alleles responsible for blood groups A and B in the JPT population than that in the YRI population (Figure S15; π= 0.023% for type A and 0.048% for type B in JPT;π= 0.115% for type A and 0.083% for type B in YRI). Notably, in this analysis, we excluded blood group O given its multiple origins.

As mechanisms of balancing selection, both overdominance and frequency-dependent selection with a rare-allele advantage are well-studied. Additionally, fluctuating selection, which involves spatio-temporal variations in selection pressure, has been proposed (Charlesworth 2006). The genetic signatures of these three mechanisms overlap, making it exceedingly challenging, often not feasible, to distinguish the underlying mechanisms from the genetic signatures alone (Spurgin and Richardson 2010; Fijarczyk and Babik 2015). In the present study, using forward simulations, we examined which mechanism, negative frequency-dependent or overdominance selection, is more appropriate to explain the maintenance of the three haplogroups and the ancestral (African-specific) haplogroup. We concluded that negative frequency-dependent selection is more likely from the functional perspective of APOBEC3 and is more parsimonious based on simulations. Under negative frequency-dependent selection, the differing patterns of genetic variation in African and non-African populations can be explained by a single combination of parameter *a* and parameter *s*. In contrast, under the overdominance model, the *a* value, which is assumed based on the ancestral group frequency in the African populations, could not explain the observation in the non-African populations, leading to the necessity of adjusting the parameter between the African and non-African populations.

The parameter *a* defines the relative fitness of the African-specific ancestral group (1 − *as*) in contrast to the relative fitness of other haplogroups (1 − *sx*). Under this frequency-dependent selection model, the haplogroup frequency *x* hardly exceeds the value of *a*. For example, if *a* = 0.2, the frequency of a haplogroup tends to be restricted to less than 0.2 in a large population. At *x* = 0.2, the haplogroup and the ancestral group possess identical relative fitness. However, at *x* > 0.2, the relative fitness of the haplogroup would fall below that of the ancestral group, resulting in the replacement of the haplogroup by the ancestral group. When *a* becomes 0.3, the frequency of each of the three haplogroups can increase up to 0.3, which is compensated by the reduced frequency of the ancestral group. Thus, the frequency and diversity of a haplogroup sensitively vary based on the different values of *a*. We envisage that difference in the value of parameter *a* reflects differences in the pathogenic viral environment.

In the forward simulation, we assumed a single nucleotide site as a target of balancing selection, with no recurrent mutation at this site. Given these assumptions, we cannot expect either haplotype turnovers or enhanced substitution rates that can occur under balancing selection (Maruyama and Nei 1981). As noted by Takahata and Nei (1990), who used the infinite-alleles model, these features are particularly relevant to the evolution of the MHC peptide binding region (Hughes and Nei 1988). In this context, although we emphasized a haplotype turnover in HI prior to the Out-of-Africa bottleneck, it does not necessarily imply the operation of independent directional selection if newly arising mutations are allowed to undergo balancing selection. We note that this may not be restricted to HI and that more general mutation models would be compatible with our sequence data. In conclusion, under different mutation schemes, joint effects of negative frequency-dependent selection and demographic changes with bottlenecks are sufficient to explain the observed pattern and level of nucleotide polymorphism in the *APOBEC3* regulatory region.

We relied on a simulation method to detect natural selection on the *APOBEC3* haplogroups, as the existing detection methods are not appropriate in the present case. Approaches for detecting positive selection, such as EHH-based methods (Sabeti et al. 2002; Voight et al. 2006), SDS (Field et al. 2016), and *F_c_* measurements (Fujito et al. 2018), use summary statistics that directly or indirectly measure the ratio of the diversity in the focal sequence group to the overall sequence diversity. This ratio is expected to be low when the focal group is under positive selection. However, the ratio cannot be very low if some remaining haplotypes have also reduced the diversity for any reason. Similarly, other summary statistics, such as *β* (Siewert and Voight 2017) and *β^(2)^* (Siewert and Voight 2020), are applied to detect clusters of alleles at similar frequencies around the site targeted by balancing selection. As such a cluster of alleles reflects the accumulation of variations in two allelic lineages maintained by balancing selection through time, the *β* and *β^(2)^* statistics gain power for low recombination rate, high mutation rate, and long-term balancing selection. However, the persistence time of the three haplogroups in the *APOBEC3* regulatory region is not long enough to accumulate a sufficiently large number of variants to be detected. Under this scenario, these statistics tools may lose power.

We obtained information that supports the existence of functional differences in the *APOBEC3* regulatory region from GTEx eQTL data and the GWAS catalog. Using GTEx eQTL data from 52 tissues, we searched for variants that might affect the transcriptional regulation of the seven *APOBEC3* genes. We found that the regulatory region harbors several tissue-specific eQTLs for the expression of all *APOBEC3* genes, except *APOBEC3H* (Figure S16, Table S6). This includes index SNPs and linked SNPs of the three haplogroups. These eQTL associations are relatively weak compared to those in the downstream region within the *APOBEC3* gene cluster (chr22:39,405,000-39,580,000; Figure S6). However, a recombination hotspot separates the downstream region from the regulatory region (Figures 1a and S16a). As a selective sweep was detected on a haplotype in *APOBEC3H* (Fujito et al., unpublished data), the downstream region appears to be under the influence of another process of evolution in terms of an arms race with viruses. Therefore, the regulatory region appears to be the only region where natural selection can act solely on the transcriptional regulation of *APOBEC3* genes. Additionally, we searched the GWAS catalog for variants in the *APOBEC3* regulatory region. We found that rs5995653, which is linked to HII, is associated with asthma exacerbation in admixed children (Hispanics/Latinos and African-Americans) treated with inhaled corticosteroids (*p* = 5 ×10^-6^; OR = 1.52); this association is also found in Europeans (*p* = 7.52 ×10^-3^) (Hernandez-Pacheco et al. 2019). These results suggest that HII affects the overall immune state.

Owing to their mutagenic characteristics, APOBEC3s can potentially harm host genomes as well. APOBEC3A and APOBEC3B are the primary sources of somatic mutations in many tumors (Alexandrov et al. 2013; Roberts et al. 2013; Alexandrov et al. 2020). The deletion polymorphism of *APOBEC3B* drives the strongest signal for APOBEC3B-like mutagenesis at the pan-cancer level (ICGC/TCGA Pan-Cancer Analysis of Whole Genomes Consortium 2020). Therefore, we examined the possibility that positive selection is triggered by this deletion. However, we found no evidence of positive selection specifically targeting the *APOBEC3B* deletion despite its functional impact. This finding suggests that the deletion confers little to no selective advantage during evolution and that its increased frequency may result simply from genetic hitchhiking of tightly linked HI. Within HI, sequences linked to the deletion may have conferred greater advantages compared to other HI haplotypes. However, the reduction in sequence diversity caused by additional positive selection acting on the deletion is likely too small to be detected. This is because sequence diversity in HI is already extremely low due to the effects of population bottlenecks and haplotype turnover. However, even in such a case, the deletion may need to be linked to HI to have evolved, because when linked to other sequences, it does not exhibit any sign of expansion (Figure S14).

Several issues should be addressed in future studies. First, more precise knowledge about regulatory functions that confer a selective advantage to each haplogroup needs to be elucidated (i.e., which *APOBEC3* gene is upregulated or downregulated as the core function of each haplogroup). This was challenging to determine in the present study as research on APOBEC3s is still ongoing, and specific immune cell types where the expression of each A3 is critical for the anti-viral activities are not yet known. We utilized the publicly available eQTL data to examine the functional differences among the haplogroups; however, the existing eQTL data are from whole-blood samples, and data for each immune cell type are not available. Second, we could not identify viral targets of each haplogroup. Further elucidation of the functions of each APOBEC3, especially single-cell eQTL analysis of immune cells, might help link haplogroups to its target viruses. This will help to clarify the detailed selective advantages of each haplogroup, enabling evaluation of the susceptibility of individuals to viral infections.

## Conclusion

We have demonstrated that the “multi-star” genealogy observed in the *APOBEC3* regulatory region in 24 TGP populations is inconsistent with neutral evolution based on coalescence simulations and genomic data. Most likely, the genealogy has been shaped by the combined effects of negative frequency-dependent selection and a severe reduction in effective population sizes, accompanied by the Out-of-Africa dispersal of modern humans. A relatively recent haplotype turnover within haplogroup I is also consistent with this hypothesis under certain mutation models. The action of such selection in the evolutionary conserved regulatory region underscores its functional importance in regulating the expression of the seven linked *APOBEC3* genes.

## Data availability

We utilized whole genome sequence data sets that are publicly available from the 1000 Genomes Project Phase 3 (1000 Genomes Project Consortium et al. 2015) and the HGSVC (Ebert et al. 2021).

## Acknowledgments

We thank Dr. Naoyuki Takahata for the useful discussions and critical comments, which greatly improved this study. We also thank Dr. Toshiyuki Hayakawa and Dr. Kohta Yoshida for their helpful discussions and comments. We thank Dr. Naruya Saitou for critically reviewing the manuscript. Computations were partially performed on the NIG supercomputer at ROIS National Institute of Genetics. This manuscript has been edited for English by Dr. Quintin Lau.

## Funding

This study was funded by the Research Organization of Information and Systems (ROIS; https://www.rois.ac.jp), by JSPS KAKENHI [Grant Number 23K05870] and Shiseido Female

Researcher Science Grant.

## Materials and Methods

### Whole-genome sequencing (WGS) data for modern humans

We used phased WGS data from the 1000 Genomes Project Phase 3 (1000 Genomes Project Consortium et al. 2015). Samples from two populations, African Ancestry in Southeast US (ASW) and African Caribbean in Barbados (ACB), were excluded owing to possible admixture. The remaining 4694 samples from 24 populations were used. For African populations, 216 samples for YRI (Yoruba in Ibadan, Nigeria), 198 for LWK (Luhya in Webuye, Kenya), 226 for GWD (Gambian in Western divisions in the Gambia), 170 for MSL (Mende in Sierra Leone), and 198 for ESN (Esan in Nigeria) were included. For non-African populations, 198 samples for CEU (Utah residents with Northern and Western European ancestry), 214 for TSI (Toscani in Italy), 198 for FIN (Finnish in Finland), 182 for GBR (British in England and Scotland), and 214 for IBS (Iberian population in Spain) among Europeans (EUR); 206 samples for GIH (Gujarati Indian from Houston, Texas, USA), 192 for PJL (Punjabi from Lahore, Pakistan), 172 for BEB (Bengali from Bangladesh), 204 for STU (Sri Lankan Tamil from the UK), and 204 for ITU (Indian Telugu from the UK) among South Asians (SAS); 206 samples for CHB (Han Chinese in Beijing, China), 208 for JPT (Japanese in Tokyo, Japan), 210 for CHS (Southern Han Chinese, China), 186 for CDX (Chinese Dai in Xishuangbanna, China), and 198 for KHV (Kinh in Ho Chi Minh City, Vietnam) among East Asians (EAS); and 128 samples for MXL (Mexican ancestry in Los Angeles, California), 208 for PUR (Puerto Rican in Puerto Rico), 188 for CLM (Colombian in Medellin, Colombia), and 170 for PEL (Peruvian in Lima, Peru) for Americans (AMR) were included. Haplotypes in the *APOBEC3* regulatory region were constructed by including SNPs on chr22: 39388993 to 39405510 (GRCh37). After removing indels using VCFtools (Danecek et al. 2011), 588 SNPs remained in the region. For time estimation, we obtained ancestral allele information from the 1000 Genomes Project Phase 3 and used 584 biallelic SNPs with ancestral allele information. The 16-kb *APOBEC3* regulatory region excluded the APOBEC3B deletion (as well as identical 350-bp sequences flanking both ends of the deletion) (Figure S2). Moreover, we checked the sequence similarity within the 200-kb *APOBEC3* cluster using a dot plot (Figure S2) (Schwartz et al. 2000). We observed a cluster of short fragments (22–1668 bp) with high sequence similarity (∼98%) between the upstream and downstream areas of the deletion. We confirmed that this cluster of sequences was not included in the 16-kb regulatory region analyzed in this study (Figure S2). We also validated read mapping accuracy using different data sets (Supplementary text).

We used a deletion polymorphism, “YL_CN_STU_4456,” in the 1000 Genomes Phase3 structural variant data as the linkage information of the APOBEC3B deletion (Supplementary text). We retrieved the regulatory features from the Ensemble webpage (“homo_sapiens.GRCh38.Regulatory_Build.regulatory_features.20220201.gff”) and subsequently lifted them to the GRCh37 reference genome using the UCSC webpage. To confirm our results, we utilized WGS data from the HGSVC (Ebert et al. 2021), which are based on long-read PacBio WGS data from 32 diverse human samples, and high-coverage (30X) WGS data from the 1000 Genome Project resource (Byrska-Bishop et al. 2022), which include data from 3202 samples (Supplementary text).

### Definition of the haplogroups

We defined three haplogroups based on the following criteria: a frequency of *f* > 0.15, an average nucleotide difference per site across all pairs between haplogroups of *d* > 0.1%, and nucleotide diversity within each haplogroup of π < 0.01% in non-African populations. In defining these haplogroups, we excluded American populations, owing to known admixture.

### Data for ancient hominins and chimpanzees

We retrieved WGS data for ancient hominins from http://cdna.eva.mpg.de/neandertal/Vindija/. We used data from the Altai Neanderthal and Denisova Pinky for the phylogenetic and network analyses. We obtained sequence data for chimpanzees (NC_036901.1) and orangutans (Susie_PABv2) from the NCBI and the Ensemble webpages, respectively.

### Phylogenetic analysis

We constructed Neighbor-joining trees using MEGA 11 software (Saitou and Nei 1987; Tamura et al. 2021) based on haplotype sequences in the 16-kb region. We computed evolutionary distances using the number of differences method. We considered data for the JPT, CEU, PJL, PUR, YRI, and LWK populations as representative of the East Asian, European, South Asian, American, and African populations, respectively.

### Identification of haplogroups and recombinants

Based on the topology of the NJ trees, we defined one of the SNPs on the branch that led to each of the haplogroups as an index SNP: rs4303815 for H I, rs35043274 for HII, and rs111468681 for HIII.

We identified and removed recombinants from the time estimation and haplogroup diversity measurements (Table S4). We calculated the number of nucleotide differences for all sequence pairs within a haplogroup to identify the recombinants. We then plotted their distributions and identified outlier pairs. Sequences that caused outliers were regarded as recombinant and removed from the analyses.

### Estimation of divergence time among the haplogroups

We estimated the divergence time between each haplogroup and other sequences using the number of nucleotide differences. Specifically, based on the phylogenetic tree using sequences from African populations (Figure S4a), we grouped four groups of sequences that do not belong to any of the three haplogroups in addition to three haplogroups. We calculate the average number of nucleotide differences from all pairs among the seven sequence groups in total. Using these calculated numbers of differences, we estimated the divergence time of the three haplogroups using the mutation rate of 0.5 × 10^-9^/site/year.

### Frequency trajectories over time

We retrieved genotype data from the Allen Ancient Genome Resource (Mallick et al. 2023; Anon) (version 52.2, downloaded from https://reich.hms.harvard.edu/allen-ancient-dna-resource-aadr-downloadable-genotypes-present-day-and-ancient-dna-data). We also used genotypes of samples from Europe (4624 samples) and East Asia (898 samples) from 100– 40,000 years ago. As the index SNPs for HI, HII, and HIII were not included in these data, we utilized rs2142833, rs9607601, and rs11089916, respectively, as tag SNPs.

### Estimation of the time of expansion of the haplogroups

We estimated the time of expansion for each haplogroup by measuring intra-allelic diversity using two different methods. The first method is estimating the mean coalescence time within a haplotype group. We calculated the average number of nucleotide differences across all sequence pairs within a haplotype; rescaled the average with the length of the sequences to obtain *π* values (per site nucleotide diversity); and then converted the *π* values to years with mutation rate (using *π* = 2*t*μ, where *t* is the time since the common ancestor of the sequences in a haplogroup and μ is the mutation rate per site per year), assuming that the mutation rate was 0.5 × 10^-9^/site/year. Standard error estimates were obtained by a bootstrap procedure using MEGA 11.

The second method involved calculating the height of the genealogy of each haplogroup (Satta et al. 2019). Based on the ancestral allele information, we identified sites where the derived allele was segregating in a haplogroup of interest, and the ancestral allele was fixed in populations outside of it. Using these sites, we counted the number of derived alleles accumulated within a haplogroup divided by the number of sequences to obtain the height of the haplogroup, which was converted to years, as described above. We calculated the variance in height using the formula described by Satta et al. (2019). For the calculation of the tree heights, we removed the sites at which *i* (*k* − 2 ≤ *i* < *k*, where *k* is the size of the haplotype group) derived alleles were identified within the haplotype of interest, but no derived alleles were present outside of it. This is because these sites likely result from recombination, which introduces a limited number of ancestral alleles into the haplotype of interest.

### Inference of selection

To demonstrate the rarity of the phylogenetic structure with three large haplogroups, we searched for a phylogenetic structure with three distinct haplogroups, each of which was comparable to or surpassed the observed structure in haplogroup size (i.e., the number of chromosomes) in both simulated data under neutrality and empirical data. We performed simulations using ms (Hudson 2002) under the two demographic models proposed for East Asians, Europeans, and Africans, which were subsequently referred to as the SFD and HFD. SFD is based on a model proposed by Schaffner et al. (Schaffner et al. 2005). HFD includes an additional population bottleneck around 900,000 years ago, which was recently identified by Hu et al. (Hu et al. 2023). We mapped a specified (observed) number of segregating sites onto a coalescent tree. The command lines used for this process are provided in Supplementary Table S9. For the empirical data, we used a window-based approach (16-kb windows with a step size of 1 kb) with the 1000 Genome data. We screened for an SNP that matched the comparable frequency observed for HI from the 5′-to 3′-ends of both simulated and empirical data, which was then defined as an index SNP of HI. Next, we searched for SNP with the exact frequency observed for HII and derived alleles only on non-HI chromosomes. The closest SNP to the 5′-end was defined as an index SNP for HII. Similarly, the index SNP for HIII was determined using the same approach.

Next, we assessed the frequency with which the observed pattern of low intra-haplogroup diversity and high inter-haplogroup divergence co-occurred in the results of neutral simulations. We retrieved 10,000 replications from the simulation described above, whereby the three allelic groups exhibited the same size as the observations (e.g., 103, 76, and 19 chromosomes for HI, HII, and HIII in JPT, respectively), and calculated nucleotide differences within and between clades. Subsequently, we investigated the rarity of simultaneously observing low diversity within each of the haplogroups and high divergence between the haplogroups under neutrality (i.e. reject neutrality if the combination is less than 1%). The comparison was performed jointly for HI, HII, and HIII.

### Negative frequency-dependent selection model

We used Monte Carlo forward simulation to explore forms of balancing selection that can consistently explain patterns of genetic diversity for African and non-African populations. We used a demographic model similar to the HSD model, which includes the Out-of-Africa bottleneck and the bottleneck in Africa around 900 KYA. We also accounted for recent population expansions by estimating the harmonic means of exponential growth, based on Gao and Keinan 2016. We assumed that the current population size for both Africans and non-Africans is 500,000 individuals. For African populations, the expansion begins from 20,000 individuals at generation 0 and reaches 500,000 individuals at generation 330. In contrast, for non-African populations, it starts from 2,000 individuals at generation 0 and grows to 500,000 individuals by generation 660. We assumed a simple simulation model where organisms in a diploid population reproduce the next generation under the Wright-Fisher model. Each organism has a chromosome of length (*L* + 1), which consists of one site under balancing selection and *L* linked neutral sites without recombination. The selected site undergoes three simultaneous mutations at the start point, defining four sequence groups that mimic three haplogroups and an ancestral sequence group. At the linked neutral sites, mutations occur at the per-site rate of μ, with reverse mutations excluded. The three haplogroups are subject to negative frequency-dependent selection, while the ancestral group possesses constant fitness. The relative fitness for haplogroups is calculated as 1 − *sx*, where *x* is the frequency of each haplogroup and *s* is the shared selection coefficient. The relative fitness for the ancestral group is defined as 1 − *as*, where *a* is a constant value. To illustrate the shits in within-haplogroup diversity (*π*) and between-haplogroup divergence and the frequency of sequence groups, we performed simulations with parameters: *L* = 160, and *s* = 0.05, 0.2 and *a* = 0.2, 0.3. In this simulation, each neutral site represents a sequence block of 100 bp, effectively mimicking the *APOBEC3* regulatory region (160 kb in length). We selected an effective population size (*N*_*e*_) and mutation rate such that the production of 4*N*_*e*_ μ corresponds to the averaged *π* over human genomes (0.1%). To reduce the simulation time, we reduced the population size and the number of generations to one-tenth of their original values, while increasing the mutation rate increased tenfold. The generation time was set to 25 years.

### Overdominance model

In the overdominance model, we modified the negative frequency-dependent selection model described above by altering the definitions of relative fitness. We assumed that all the four sequence groups are under overdominance. The relative fitness for homozygotes of each haplogroup is calculated as 1 − *s*, and as 1 − *as* for homozygotes of the ancestral group. The relative fitness for heterozygotes is assumed to be 1.

## Notes

### Competing Interest Statement

The authors have declared no competing interest.

### Summary of Updates

In the original version, we discussed the timing of haplogroup expansions and their possible association with viral pandemics. During revision, we recognized the importance of explicitly considering the effects of population bottlenecks. We therefore revised the manuscript substantially, incorporating simulation-based analyses to explore the joint effects of bottlenecks and balancing selection on intra-haplogroup diversity. Accordingly, the focus of the study shifted from viral pandemic-driven positive selection to broader evolutionary mechanisms shaping haplogroup diversity.

